# Viscoelastic Parameterization of Human Skin Cells to Characterize Material Behavior at Multiple Timescales

**DOI:** 10.1101/2021.07.09.451793

**Authors:** Cameron H. Parvini, Alexander X. Cartagena-Rivera, Santiago D. Solares

## Abstract

Countless biophysical studies have sought distinct markers in the cellular mechanical response that could be linked to morphogenesis, homeostasis, and disease. Here, a novel iterative-fitting methodology is used to investigate the viscoelastic behavior at multiple relaxation times of human skin cells under physiologically relevant conditions. Past investigations often involved parameterizing linear elastic relationships and assuming purely Hertzian contact mechanics. However, linear elastic treatment fails to capture and properly account for the rich temporal information available in datasets. We demonstrate the performance superiority of the proposed iterative viscoelastic characterization method over standard open-search approaches. Our viscoelastic measurements revealed that 2D adherent metastatic melanoma cells exhibit reduced elasticity compared to normal counterparts—melanocytes and fibroblasts, whereas are significantly less viscous than only fibroblasts over timescales spanning three orders of magnitude. Interestingly, melanocytes are stiffer than melanoma cells, while being the less viscous cells measured. The measured loss angle indicates clear differential viscoelastic responses across multiple timescales between the measured cells. We propose the use of viscoelastic properties at multiple timescales as a mechanical biomarker of diseases. Altogether, this method provides new insight into the complex viscoelastic behavior of metastatic melanoma cells relevant to better understanding cancer metastasis aggression.

## Introduction

Current state-of-the-art mechanobiological applications involve testing samples that are soft, viscous, and/or polymeric in nature [1]. Understanding the mechanical character of these materials at the nanoscale is especially important in biological studies [2, 3, 4]. Whether the goal is characterizing biophysical behavior in human lung epithelial cells [5], describing the various microrheological phases that isolated adherent cells generally exhibit [6], investigating why cancer cells apparently soften during disease progression [7], or investigating why human cardiac cells showcase both shear-force fluidization and strain-stiffening while beating (despite appearing contradictory) [8], using an appropriate analytical treatment for soft sample mechanics is critical to correctly describing complex biological processes.

Cancers are a group of diseases that possess a common characteristic of leading to the development of transformed cells exhibiting a rapid and uncontrollable burst of growth and proliferation, which in turn leads to formation of a solid tumor [9]. Metastatic cancer is an aggressive disease, characterized by its high level of mutational burden, resistance to traditional chemotherapies, and rapid metastasis dissemination [9]. To successfully metastasize, cancerous cells must migrate away from the primary tumor, invade the surrounding tissue, extravasate through the circulatory system, and intravasate through the vasculature to colonize a new tissue [10, 11]. While cancer cells can use a variety of migration strategies, metastatic cells are subjected to the stringent physical constraints of the extracellular matrix (ECM) [11, 12, 13]. Thus, to successfully pass-through micron sized gaps in the ECM, the cell actomyosin cortex and its nucleus (the largest and stiffest organelle in the cell) must be deformed to a high extent (over 50% to 80% of its original size), thus experiencing large external mechanical stresses [12, 14]. Therefore, the mechanical properties of cancerous cells, including melanomas, are critical for successful survival, migration, and colonization of tissues.

Recent work has shown that the intracellular mechanical properties are mostly controlled by the cytoskeleton, a network of intermediate filaments, microtubules, and filamentous actin [15]. Cells are often considered to be soft biomaterials that behave as complex viscoelastic fluids in terms of their response to external mechanical stresses [16, 17]. By applying Atomic Force Microscopy (AFM), cancer cells have been shown to be more deformable than their normal untransformed counterparts [7, 18], potentially due to modifications in cytoskeletal organization, possibly aiding them with upregulated migration through a structurally and mechanically complex ECM [19, 20, 21]. However, most of the current cell mechanics research has relied upon continuum mechanics models which assume cells exhibit purely elastic behavior [22]; this approach has been frequently applied when characterizing cancerous cells [7, 23, 24, 25, 26, 27, 28, 29]. While this methodology may be useful under very specific conditions, this linear elastic treatment fails to capture and properly account for the rich temporal information available in soft sample mechanical behavior datasets. For example, in applications using AFM to evaluate cell properties, the difference between assuming linear-elastic sample behavior versus more complex viscoelastic mechanics can be significant [30, 31, 32, 33, 34]. While there have been a few investigations into linear viscoelastic assumptions for soft samples [35, 36, 37], these implementations rarely account for more than two discrete stiffnesses and one retardation time. For cellular action taking place over a longer period, such as during cancer metastasis extravasation or intravasation, a single retardation time could not simultaneously account for both small- and large-timescale mechanical responses. In this case it is necessary to approximate physical action using a model that successfully delineates between timescales, such that there still remains a need to develop generalized viscoelastic models for soft cells.

Using a modified approach outlined in this paper, we build upon a previous open-search viscoelastic parameter extraction methodology [31] to parameterize biological samples with a dynamic number of retardation times, and search for new mechanical information that emerges at multiple timescales. One clear application is for use on cancerous cells, where directly tracking the effects of each viscoelastic model element could uncover new mechanical signatures that are useful for the early detection of common cancers, such as metastatic melanoma. The method presented here also reduces the computational overhead for each individual parameter fit by limiting the associated parameter space. This decreases the number of iterations necessary to obtain each unique mathematical series describing the material, leading to comparable overall parameterization times and less susceptibility to local minima versus the previous methodology. While this iterative approach still allows for further optimization, it provides a marked improvement over the open-search methodology and has been applied successfully for soft samples.

This study begins by introducing the new iterative method used to improve the accuracy of the fitting functions, showing that the performance of the new approach is far superior and more accurate than standard open-search methods. We then utilize the novel method to characterize the viscoelastic behavior at multiple retardation times of 2D adherent human metastatic melanoma cells and compare them with their normal counterparts, primary epidermal melanocytes and fibroblasts. Our viscoelastic measurements reveal that 2D adherent metastatic melanoma cells are less elastic and moderately viscous over the relaxation timescale studied when compared to melanocytes and fibroblasts, whereas fibroblasts are the stiffest and more viscous. Interestingly, melanocytes have intermediate stiffness, while they are the least viscous. Altogether, the study provides a general understanding of the complex viscoelastic behavior of living metastatic melanoma, which is relevant to better understanding metastasis and aggression.

## Results

### Improved Characterization of Cells Viscoelastic Behavior by Parameterization Approach

One shortcoming of using an open-search methodology [38] to parametrize a viscoelastic material using a discrete generalized model is the inability to pinpoint the contribution of each viscoelastic element in the model to the overall mechanical response. This is due primarily to the fact that model elements are not individually treated during the fitting process—all parameters are estimated simultaneously using a nonlinear least-squares regression approach (Fig. 1). The associated parameter space for each fit is large, which can easily lead to improper coupling of different elements. For example, in the case of a multi-element material model, the fitting algorithm could artificially increase the stiffness of one particular element to account for an improperly tuned characteristic time in another element. The algorithm would therefore skew the stiffnesses of both elements and fail to correct the error in the characteristic times, while still providing an apparently good fit to the experimental dataset. This shortcoming of the open-search method may not be critical when the user only seeks a close approximation, but can be detrimental to the quality of the final fit, and may require refinement at an additional, unnecessary cost. Aside from these associated computational penalties, the range of values which provide an adequate fit to the data can have drastically different implied mechanical properties, which in turn complicates and adds uncertainty to the analysis of the sample response. These ambiguities seem to be exacerbated in the case of soft biological samples.

**Fig. 1.**
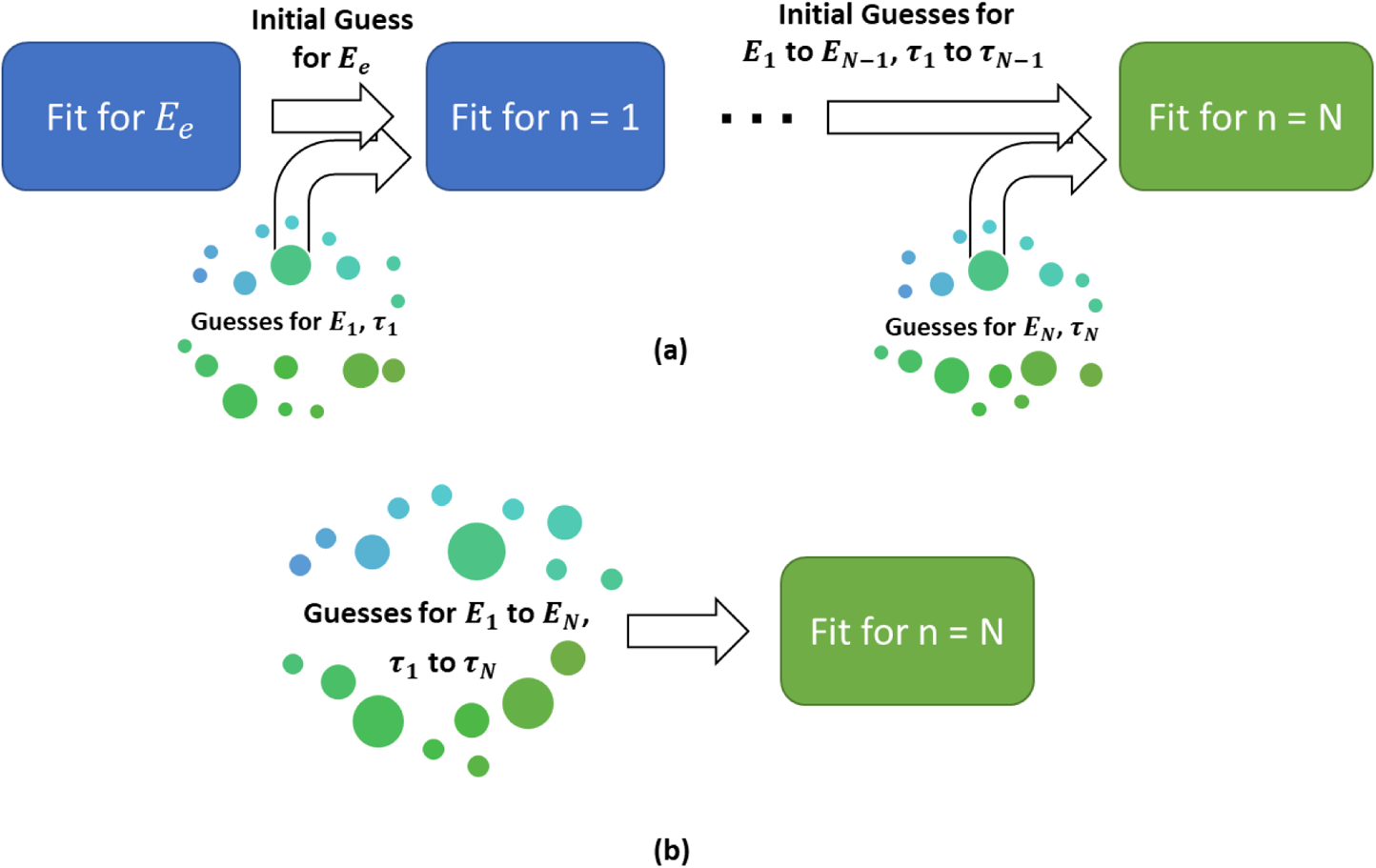
Process visualization for the Iterative-Fitting (a) and Open-Search (b) methodologies using increasingly more complex configurations within a generalized viscoelastic model (see Fig. 2f). As depicted, the Iterative-Fitting methodology involves slowly expanding the desired generalized viscoelastic model while iteratively re-parameterizing each new configuration. The “optimal” parameter set for a given configuration is used as the initial guess in the next fitting attempt. In this scheme, a random starting point is generated only for the new parameters. In the Open-Search methodology, there is no a priori information provided to the parameterization process, aside from the valid parameter bounds and a random initial guess for each parameter in the model. The variable “N” represents the N^th^ element introduced into the generalized viscoelastic model.

To evaluate the efficacy of both the open-search and iterative approaches, the elapsed time was tracked while a simulated AFM quasi-static force curve was used to parameterize a generalized viscoelastic model. The results, in addition to the fit quality as a function of the number of terms introduced, are shown in Fig. 2 for both approaches. For this evaluation case, 500 fitting iterations were run for each unique series, with up to four timescales (each requiring one separate term in the model), beginning with an initial timescale on the order of 10^−4^ seconds and using increasing timescales for each new model term.

**Fig. 2.**
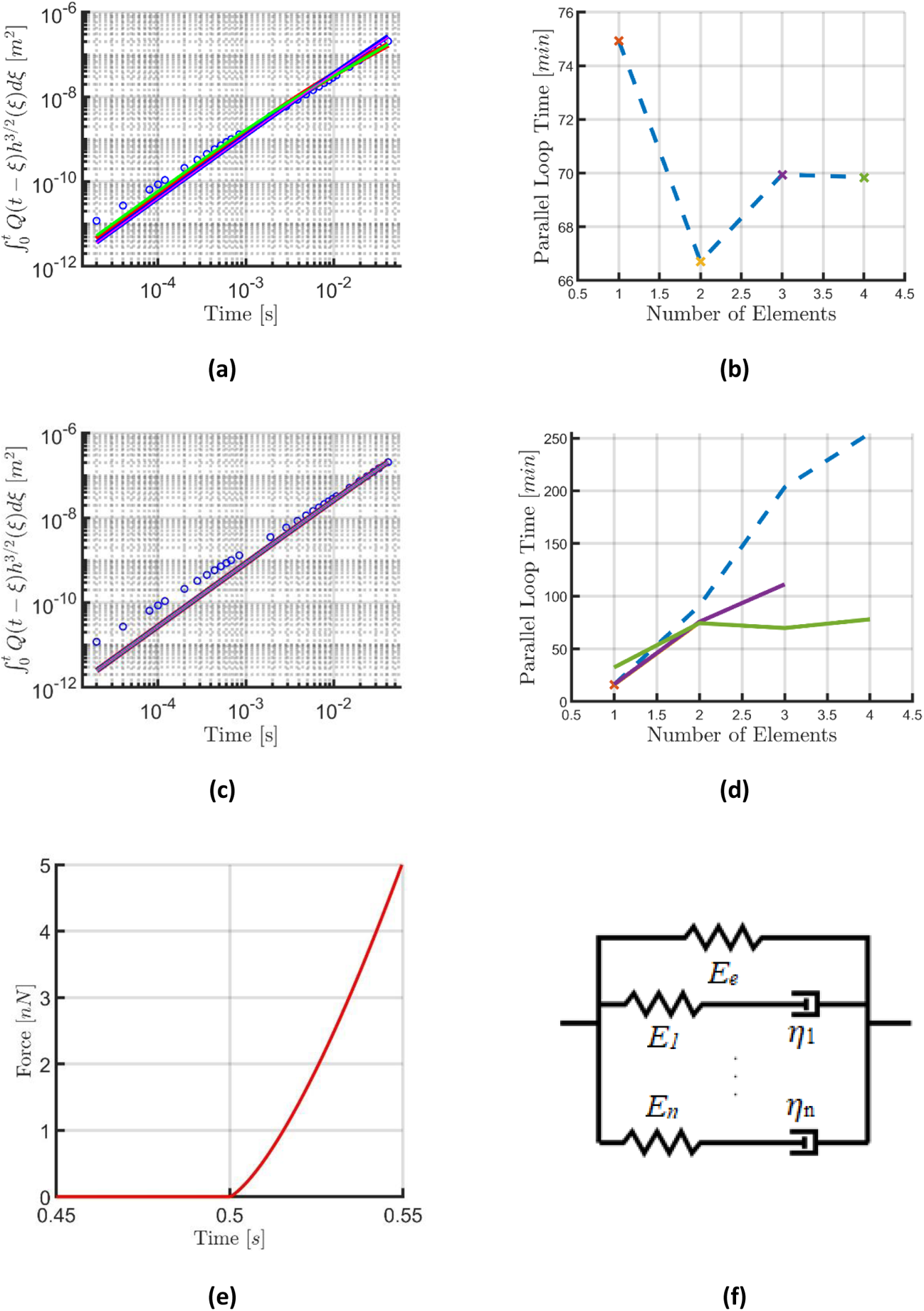
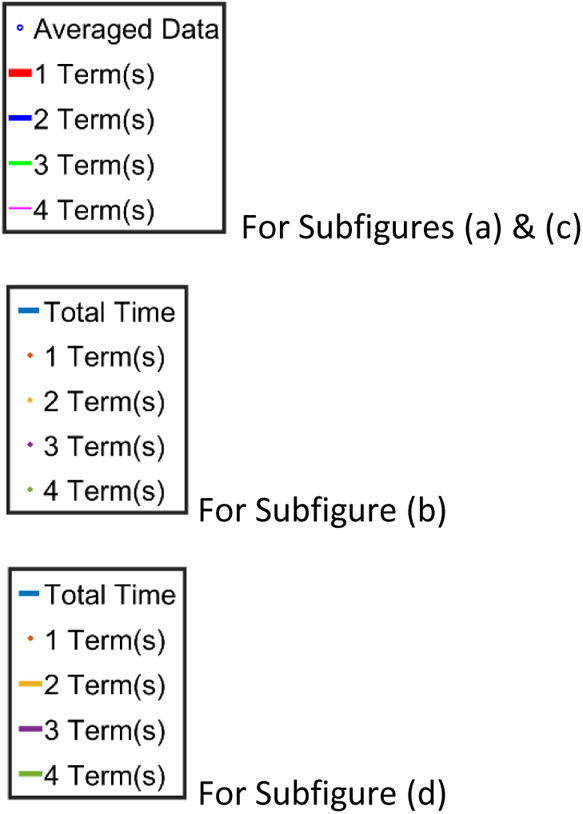
Fit performance and computational requirements for the Open Parameter Search Method (a & b), and the Iterative Term Introduction Method (c & d), in addition to the original force curve (e) generated using the Generalized Maxwell Viscoelastic Model (f) with parameters provided in the Supporting Information. The dashed blue lines in (b) and (d) represent the total time necessary to acquire a set of best-fit parameters for a model with the number of elements indicated — i.e., for 4 elements, the Iterative Term Introduction Method would require just over 250 minutes to run the fitting for terms 1-4, while the Open Parameter Search Method needs only the time indicated for the 4-term fit (approximately 70 minutes). The fitting time for the iterative approach using 1 term is lower than the corresponding fit for the Open Parameter Search Method because in the former the elastic term is first fit independently, and then fed forward to the 1-term parameterization attempt.

The elapsed fitting time (Figs. 2b and d) clearly shows that the iterative approach required a longer time to fit an equivalent number of terms when compared with the open-search approach. This is expected, since the iterative method requires separate calculations for each number of elements leading up to the desired final quantity. Although the overall time requirement was larger, the fitting time of each term in the iterative case (considered individually) was of similar order of magnitude as for the open-search case. In addition, the nearly overlapping lines in the iterative case showcase its remarkably stable performance (Fig. 2a), in contrast with the relatively small but non-monotonic variations in the open-search (Fig. 2c). This observed instability for the open-search method can lead to degraded performance for experimental data sets exhibiting significant noise contributions. Overall, when evaluating the performance and fitting time requirement, the iterative approach methodology required a moderate increase in time while providing a more repeatable performance. In addition, when the user is interested in evaluating the results as a function of the number of terms introduced (this is recommended in order to avoid “overfitting” the data, since the required number of viscoelastic model elements is not known a priori), the iterative method can provide that information faster relative to the open-search method. For example, summation of all points in Fig. 2b reveals that the total time to obtain this information for the open-search method (~280 minutes) is larger than the “Cumulative Time” in Fig. 2d (~250 minutes) for a four-term fit using the iterative method. If more elements are desired, the difference in time required to obtain this information would become increasingly different for the two approaches.

### Minimal Parameterization Terms Required for Cell Viscoelastic Characterization

The improved stability of the iterative parameter extraction methodology enables its application to samples that were previously difficult to analyze and differentiate, making it especially relevant for biological and other soft samples. To implement this approach, AFM force curves were collected from 2D adherent human metastatic melanoma cells, as well as their normal counterparts, primary epidermal melanocytes and fibroblasts at the nuclear region (Fig. S1).

The critical point of merit for a parameterized model’s performance is how closely it reproduces the normalized dataset. In this case, indentation depth and force have been experimentally measured with AFM and scaled according to one of the following equations:

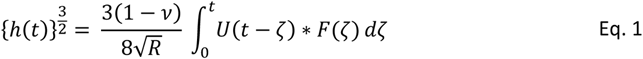

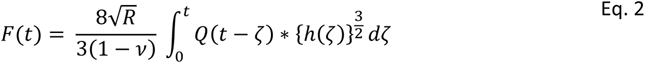

Here, the measured indentation (h) and force (F) have been convolved with the model-specific viscoelastic Relaxance (Q) or Retardance (U), and then scaled by the tip radius (R) and Poisson ratio (ν), in accordance with the well-known Lee and Radok spherical indentation framework [39]. Using each compliance- (Eq. 1) or stiffness-based (Eq. 2) model description, the convolution term has been calculated using the best-fit parameters and the repulsive (i.e., compressive force application) portion of the indentation force curve. For a full description of the analytical methods applied when deriving the material Relaxance (Q) or Retardance (U), see the Supporting Information, which also provides detailed guidelines for the choice of a viscoelastic material model. The optimal fit quality is visualized here for each unique cell type within the stiffness-based Generalized Maxwell Model description (Eq. 2) [40]. Calculations were also performed using the compliance-based Generalized Kelvin-Voigt Model (Eq. 1) which yielded similar results. This behavior is expected since both models are physically equivalent [40].

As is evident from Fig. 3c-e, the introduction of additional terms into the model does not always significantly alter the resulting action integral representation of the normalized datasets—in each case, the observation of subtle improvements at the longer timescales was the primary indicator of the optimal model’s configuration within the chosen viscoelastic model representation. This highlights the need to iteratively introduce elements: the qualitative effects of each element must be well understood and must provide clear benefits over simpler configurations in the frequency range of interest in order to be useful. In the cases shown, two terms were used to fit the melanocyte dataset (Fig. 3c), three elements were necessary to fit the melanoma dataset (Fig. 3d), and two terms were necessary for the fibroblast dataset (Fig. 3e). Notice that adding a second term to the melanocyte model improved the fit, however including additional terms beyond three only served to degrade the fit quality. Clearly, inclusion of a large number of elements in the model is not always beneficial.

**Fig. 3.**
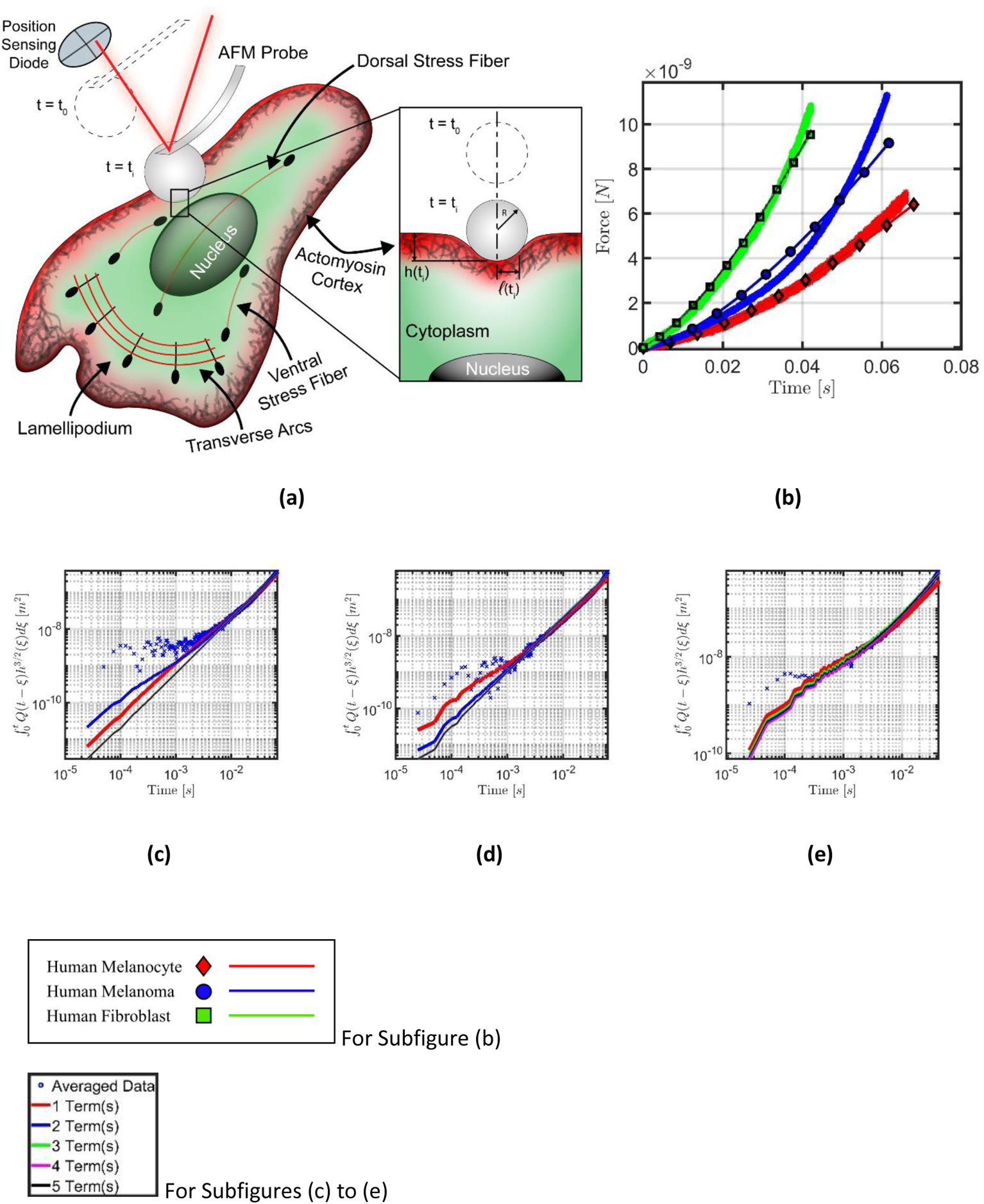
Model performance vs. action integral data for 2D adherent human skin cell lines. The experiment configuration is rendered in (a). Note that h(t_i_) is the final indentation depth and l(t_i_) the final contact radius. The force curves reconstructed using the optimal parameter sets for each adherent cell type are provided in (b), where the thick colored lines represent the averaged dataset and the thinner, marked lines represent the optimal model estimations. For subfigure the melanocyte model was fit using 2 viscoelastic elements, the melanoma model using 3 elements, and the fibroblast model also used 2 elements. The experimental data collected through the AFM force curves has also been visualized in terms of an action integral in log-spaced form as scattered markers (“Averaged Data”), and the corresponding colored lines show the action integral predicted by the model’s approximation of that dataset for a varying number of terms for melanocytes (c), melanoma (d), and fibroblasts (e), respectively. These subfigures are the basis upon which the “optimal” parameter sets are determined; the lowest number of terms that most accurately represents the input data has been selected in each case. In some cases, the first order of magnitude was difficult to approximate (as is evident in (c-e), especially for melanoma), due in part to data acquisition frequency limitations and the cost of performing numerical convolutions. This figure showcases results for the Generalized Maxwell Model in the Lee and Radok framework. Panels (b)-(e) show the results for a single representative force curve taken at the nuclear region of the cell; this process was repeated for between 70-193 AFM force curves from each cell type.

Overall, the fit to the normalized AFM observables (Fig. 3b) indicated that the chosen parameters can be used to generate satisfactory representations of the material viscoelastic behavior for most timescales, within the leading order of magnitude. Nevertheless, it is important to point out that the linear viscoelastic model chosen was unable to fully reproduce the AFM force curves, in particular the apparent stiffening behavior observed at longer timescales (larger indentation values) in Fig. 3b. In all three cases the experimental force curves initially rise slowly, and afterwards rise more sharply, with a functional dependence that corresponds to a slope that is greater than the slope allowed by the chosen viscoelastic model. Specifically, in an elastic spherical indentation experiment the force is proportional to the indentation raised to the power 3/2. For a viscoelastic material represented by the Generalized Maxwell model, the exponent on the indentation will be smaller than 3/2, since the sample experiences relaxation, and this exponent can only decrease (not increase) at longer timescales. Thus, while the chosen model offers a significant improvement over the state of the art, especially over approaches relying on purely elastic approximations, the complexity in the mechanical response of biological materials may require more elaborate mechanical models, depending on the level of accuracy sought in representing their material behavior. This is due to the fact that such systems are generally heterogeneous and nonlinear, and are often characterized using geometries which are not idealized (e.g., our systems are not infinite, perfectly flat surfaces),

### Multiple Timescales Viscoelastic Characterization of 2D Adherent Normal and Cancerous Human Skin Cells

The viscoelastic harmonic functions, storage and loss modulus, for 2D adherent normal and cancerous human skin cells are plotted in Fig. 4. These results are the figure of merit for viscoelastic analysis, which indicate how elastic or viscous the tip-sample interaction was as a function of deformation frequency (or timescale, which is the inverse of frequency). A large storage modulus indicates a strongly elastic action (increased stiffness), and similarly, a large loss modulus indicates a strongly viscous action (increased viscosity) at the corresponding frequency.

**Fig. 4.**
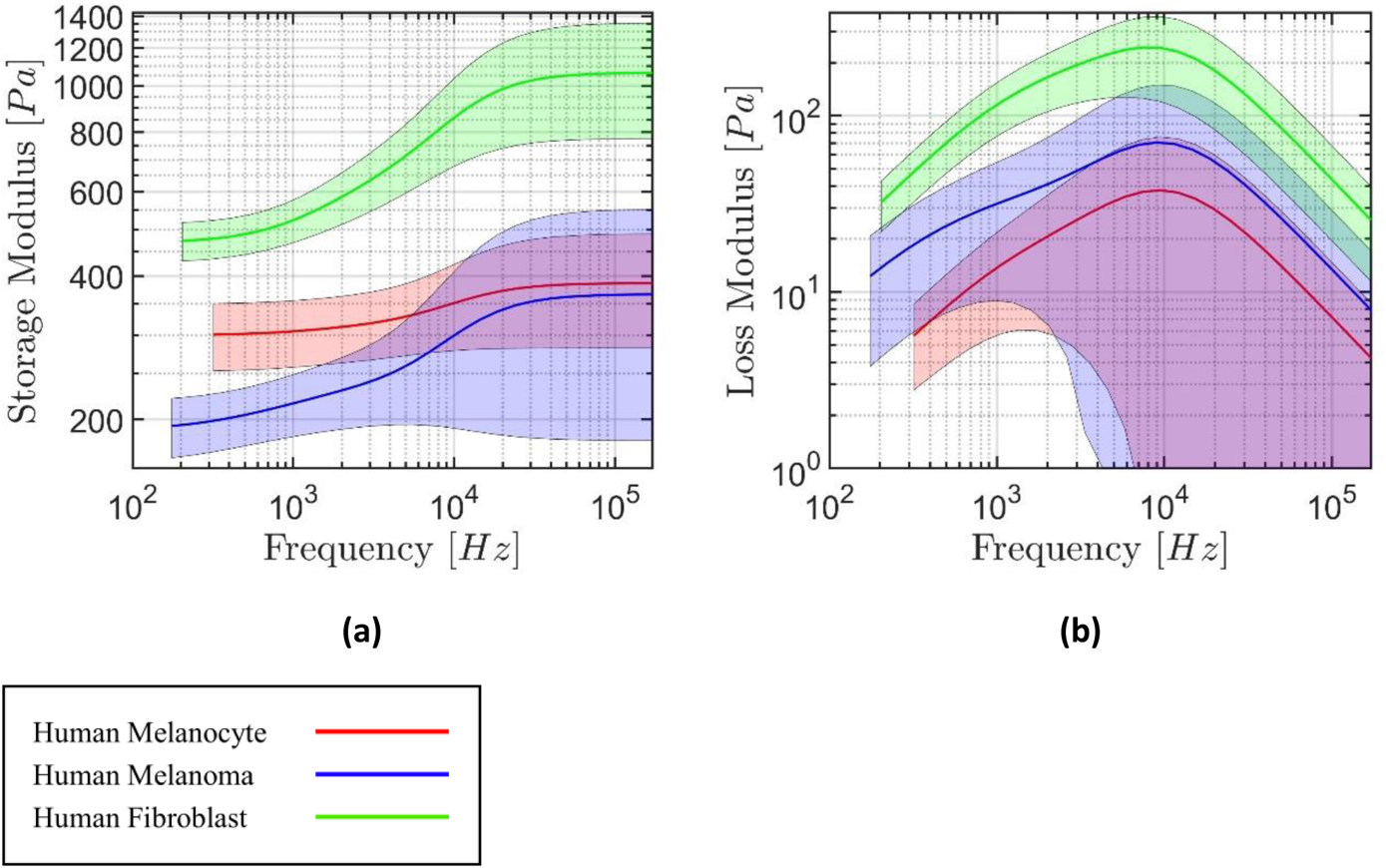
Storage (a) and Loss (b) modulus calculated from the parameterized Generalized Maxwell Model for 2D adherent normal and cancerous human skin cells. These viscoelastic harmonic quantities are calculated using the optimal parameter set obtained from analyzing an average of 70, 71, and 193 force curves from the melanoma, melanocyte, and fibroblast cell conditions, respectively. In total, the averaged datasets contain curves collected from 13 unique melanoma cells, 12 unique melanocytes, and 33 unique fibroblasts. Note that the number of terms in the fitted viscoelastic model series determines the number of distinct features present in the plots. The observed confidence intervals are shaded in matching colors for each model; the method for determining these bounds is outlined in the Materials and Methods section, and is based on the optimal parameter sets obtained using every individual curve from each cell type.

In general, all three cell types analyzed appear soft, showing storage moduli on the order of 10^2^ to 10^3^ Pa. The elastic response to indentation noticeably decreases below 10 kHz for all three cell types. The largest relative change occurs for the melanoma and fibroblast datasets, which exhibit a nearly 50% reduction in elastic action between the timescales under study. Both the melanocytes and fibroblasts exhibited relatively tight confidence intervals for the storage modulus. By contrast, the melanoma dataset features a remarkably wide confidence interval at the higher frequencies. The largest viscous action was observed for the fibroblast dataset, peaking near the onset frequency of elastic stiffening (~ 10 kHz). The melanoma dataset showcased moderate viscous action, which similarly peaked near 10 kHz, but also showcased a distinct feature for frequencies below 10 kHz; the decrease in viscous action appears to temporarily diminish at a lower rate between 500 Hz and 1 kHz before declining more sharply with decreasing frequency (longer timescales). Lastly, the melanocyte dataset exhibited the lowest viscous action, peaking once again near the onset of elastic stiffening, with loss modulus values that are approximately one order of magnitude below those of its storage modulus (~10^1^ Pa as compared to ~10^2^ Pa, respectively). Interestingly, despite the apparently good fit to the force curves in Fig. 3b for melanoma and melanocytes, the confidence intervals are relatively large for the corresponding viscoelastic harmonic quantities plotted in Fig. 4 (blue and red traces), especially for the loss modulus, which may be due to factors such as variability in the overall viscoelastic behavior of the sample, noise, or mechanical nonlinearities not captured by the model. In fact, the metastatic melanoma cells used in this study are highly dynamic and migratory, therefore possibly the higher turnover of the cytoskeleton structures and translocation of the melanoma cells would impact the viscoelastic harmonic quantities. It is noteworthy that the averaged indentation response for every cell line exhibits a clear peak in viscous action at nearly the same frequency, despite representing very different force curves in Fig. 3b.

In addition to the results presented in Fig. 4, the traditional Hertzian spherical contact model was parameterized to provide a pseudo-elastic (Young’s) modulus for each cell type, yielding the results shown in Fig. 5 for the same datasets visualized in Figs. 3 and 4. The median Young’s modulus values are appreciably larger than the moduli plotted in Fig. 4, with the Hertz model Young’s modulus values ranging from ~600 Pa to 1.2 kPa and the viscoelastic storage modulus values ranging from ~200 Pa to 1.1k Pa. Nevertheless, the modulus values for both models were within the same order of magnitude and ranked the cell type stiffnesses in the same order. Similar to the viscoelastic model fits, the Hertzian model fits did not always properly reproduce the curvature of the experimental force curves (Fig. 5b), both over- and underestimating the forces for the short and intermediate timescales (0.1 to 10 milliseconds), especially for the melanoma. In fact, the fit quality for the chosen force curves obtained with the Hertzian model seems to be comparable to the fit quality obtained with the viscoelastic models, which partly explains why the simpler elastic treatment is often assumed to be satisfactory. However, the critical shortcoming of the Hertzian approach is that it lacks the proper physics, reducing the material response to a single number that is independent of deformation rate, which makes it unable to describe the material viscous response. Generalized viscoelastic models, by contrast, incorporate the proper physics and provide a richer perspective from which material behavior at different deformation timescales can be inferred.

**Fig. 5.**
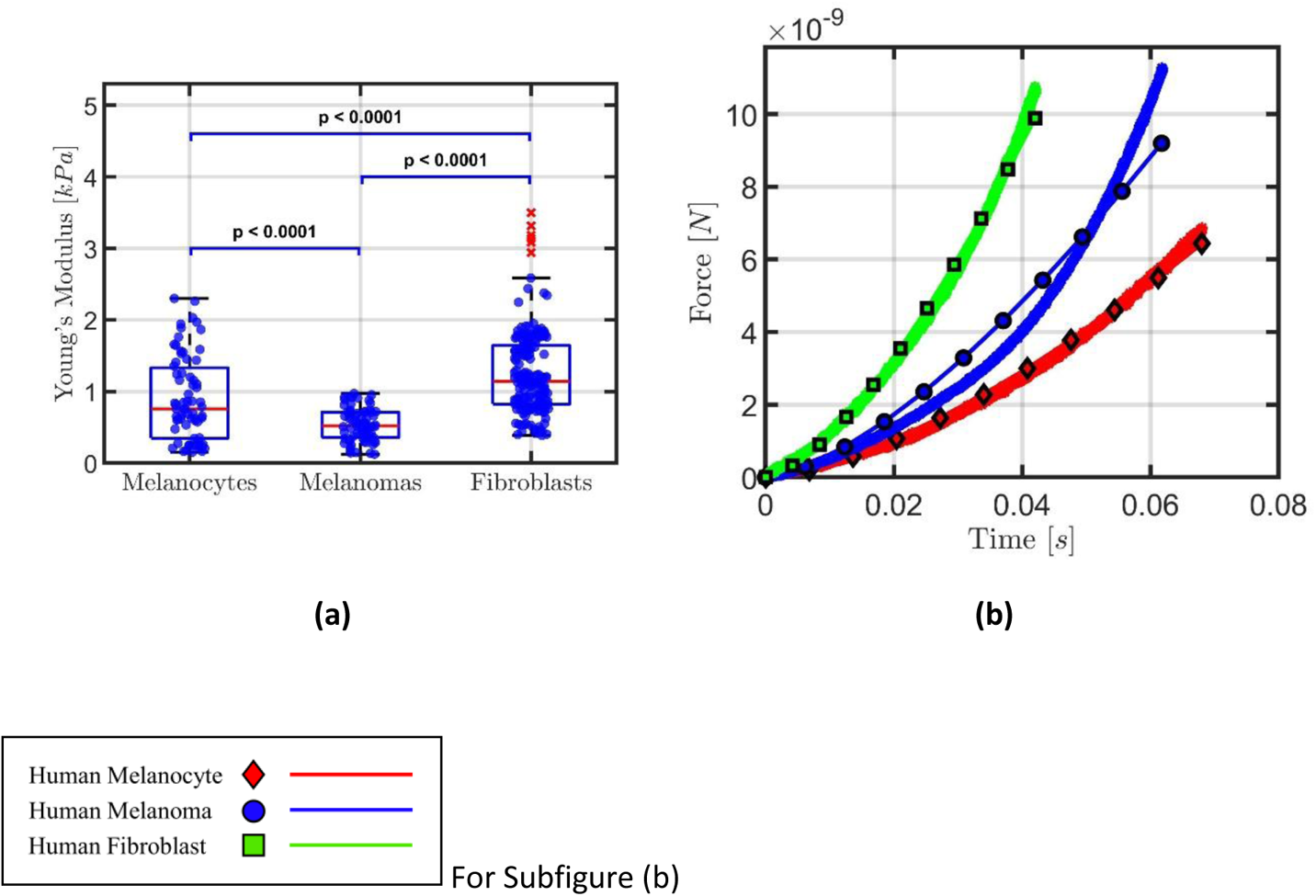
Pseudo-Elastic (Young’s) modulus distribution (a) and examples illustrating the fit quality for the Pseudo-Elastic model (b) for the adherent human skin cell lines visualized in Fig. 3. This plot provides some insight regarding the overall stiffness for each cell type, specifically that the melanoma cells appear softer than their healthy counterparts by a noticeable margin. The red markers indicate “outlier” values, which are determined to be more than 1.5 times the interquartile range away from the upper and lower bounds of the boxes. The Fibroblasts showcase the only outliers by this definition, which can be reasonably expected because significantly more curves were acquired for that cell line. The number of cells analyzed per cell type is n=12 melanocytes, n=13 melanoma cells, and n=33 fibroblasts. Data are represented as mean ± deviation with significant difference between cell types determined by unpaired two-tailed Student’s t-test with Welch’s correction indicated as, *P<0.05.

In analyzing viscoelastic behaviors, it is helpful to consider not only the magnitude of the storage and loss moduli, but also the implied loss angle. This quantity, which varies between 0° and 90°, is the inverse tangent of the ratio of loss modulus to storage modulus and provides information on the *relative* magnitude of viscous to elastic action present within the dataset at different timescales. A higher loss angle indicates proportionally greater viscous action, whereas a smaller value indicates proportionally greater elastic action. The loss angle for the three distributions shown in Fig. 4 has been plotted in Fig. 6 together with results for human lung epithelial cells from the literature [5]. The lung epithelial cell results are based on fitting the mechanical behavior measured with AFM to a Power-Law Rheology (PLR) model, for which the response of the material must follow a specific monotonic trend defined by the power-law parameters. This is in contrast to our use of generalized models, which do not constrain the response at a given timescale to exhibit any specific trend with respect to the response at other timescales. From the loss angle plots in Fig. 6, it is evident that the melanocyte dataset (plotted on the left-side axis) exhibited a remarkably low loss angle for the range of timescales studied. The averaged melanocytes loss angle peaked at just under 0.3°, and at a slightly higher frequency when compared to the fibroblasts (~5 kHz, plotted on the right-side axis). The melanoma (also plotted on the right-side axis) showed a moderate loss angle, peaking at approximately 4.5° and remaining relatively constant for frequencies below 5 kHz, in contrast to its healthy counterparts, for which the loss angle dropped more sharply at low frequencies relative to its peak value. This indicates a uniquely heterogeneous response for melanoma, which could not be characterized using a purely elastic treatment like the Hertzian model analysis of Fig. 5. The averaged melanoma response also showcases a distinct bimodal curvature, further differentiating it from melanocytes and fibroblasts. Lastly, the fibroblasts showcased the largest viscous response peak for all cell lines near 3 kHz and 7° but fell below the average melanoma response at low frequencies. The shapes of the curves for both healthy cell lines were similar to one another, although the magnitudes of their responses were separated by approximately a factor of 10.

**Fig. 6.**
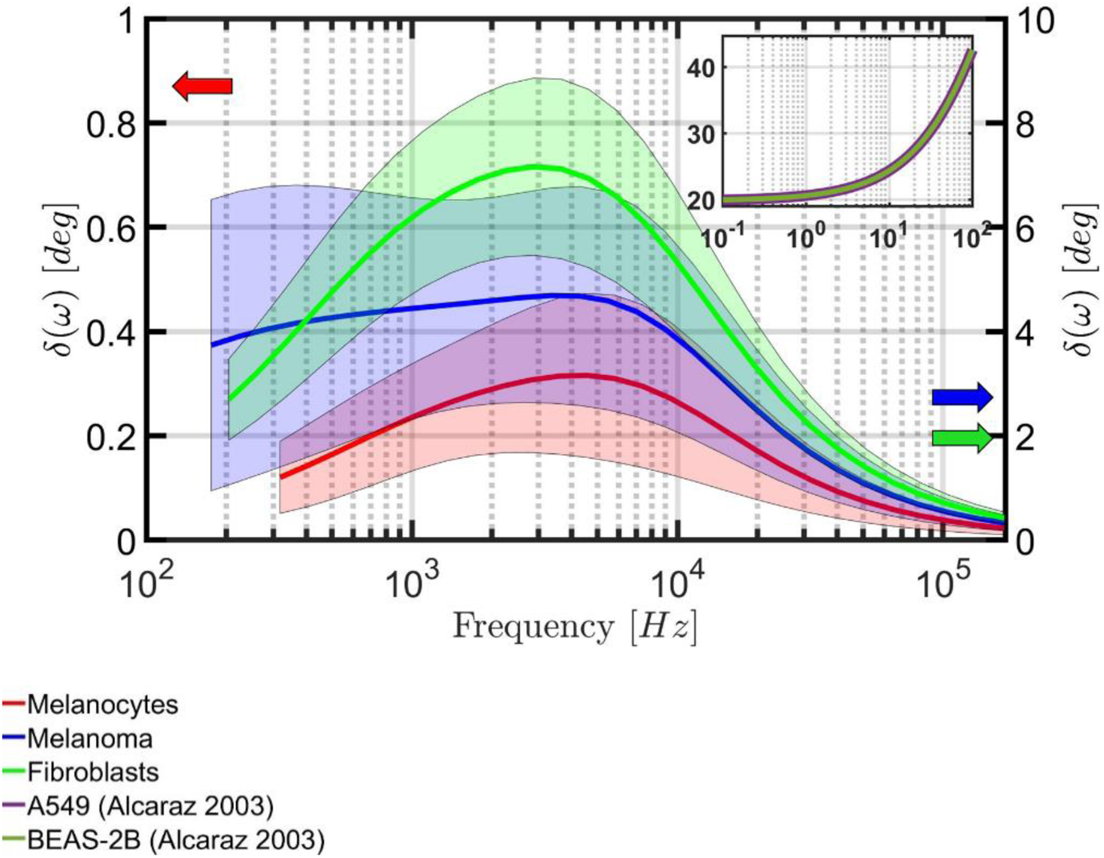
Loss Angle Comparison of the Generalized Maxwell Viscoelastic Model and Power Law Rheology (PLR) Models for Various Adherent Human Skin Cell Types. The generalized viscoelastic models showcase non-monotonic behavior for all human skin cell lines across a wide frequency range when compared with the relatively smooth and monotonic PLR model prediction for human lung epithelial cells. Observed confidence intervals are shaded in matching colors for each model. When comparing all three skin cell types, the melanocyte and fibroblast models showed noticeably tighter confidence intervals than the melanoma. As with Fig. 4, these loss angle estimates were obtained by analyzing an average of 70, 71, and 193 force curves from the melanoma, melanocyte, and fibroblast cell conditions, respectively. In total, the averaged datasets contain curves collected from 13 unique melanoma cells, 12 unique melanocytes, and 33 unique fibroblasts.

The proportion of viscous to elastic action for the three cell types studied fell far below that of the human lung epithelial cell fits to the PLR model, which exhibited monotonically increasing loss angles above 20° for a range of test frequencies below those acquired here. Clearly, since the cell types are different, it is not expected that the lung cell loss angle predictions should coincide with those of the adherent human skin cells. Nevertheless, it is insightful to compare the magnitude of the loss and storage moduli for different cell types. For both the A549 and BEAS-2B lung epithelial cell samples, the storage modulus increased nearly linearly with frequency from approximately 400 Pa to approximately 1.1 kPa, whereas the loss modulus increased with frequency, from just over 100 Pa to approximately 1.1 kPa in a manner similar to that depicted in the inset of Fig. 6. Both lung epithelial cell lines displayed nearly identical response, with loss and storage moduli on a similar order of magnitude across the frequency spectrum, but with only the loss modulus exhibiting nonlinear changes with frequency. This stands in contrast with our results presented here, for which the loss and storage moduli do not always monotonically increase or decrease, and in fact can show specific regions of the frequency spectrum where viscous action peaks. As expected, the PLR results exhibit relatively few features in the data, aside from the nonlinear, monotonic increase in viscous action.

## Discussion

As depicted in Fig. 4, all measured human skin cell types appear relatively soft when deformed with a spherical probe at their nuclear region, ranging from approximately 200 Pa to 1200 Pa in total stiffness, but their storage and loss moduli vary widely in both their absolute values and in the proportion of viscous to elastic action (Fig. 6). Furthermore, the 2D adherent metastatic melanoma sample exhibited a uniquely wide confidence interval for both moduli. All cell types exhibited their strongest viscous action near 10 kHz (Fig. 4b), and mostly displayed decreasing viscous action below and above those frequencies, with the melanoma and fibroblast datasets showing larger loss moduli than melanocytes (for the averaged response of all curves). These observations suggest that analysis of the frequency-dependent viscoelastic behavior of human skin cells could be used to unambiguously differentiate cancerous cells from their normal counterparts.

For all cell types, the proposed iterative methodology delivered a satisfactory fit of the force curves observed in the AFM experiments. However, there still remained features which were not represented well, as discussed above for all three cell types. It is important to realize that the inability of the chosen mechanical model to reproduce such features indicates that the corresponding viscoelastic behaviors are excluded from the implied viscoelastic properties (Figs. 4 and 6). Furthermore, in the application of the methodology one realizes that there exist multiple parameter sets which can reproduce the normalized data to a similar degree of accuracy. This means that iteratively introducing new terms into the model, while limiting the parameter space to some extent, does not sufficiently isolate the global-optimum parameters representing the data. Since each of these parameter sets also corresponds to a different viscoelastic harmonic response, there remains uncertainty in the extracted viscoelastic properties. This is visualized in the exceptionally wide confidence intervals for the melanoma data. While the wide confidence intervals can be seen as a shortcoming of the method or the model, they do provide additional physical insight that complements the “best fit” result. For example, inspection of the melanoma storage modulus plot (blue line) in Fig. 4a suggests that the elastic response of the material is much less than that of the melanocytes. However, consideration of the confidence interval suggests that the material stiffness could vary appreciably over the frequency range and come quite close to the melanocyte response for high frequencies. Similarly, inspection of the corresponding loss modulus plot in Fig. 4b suggests that a softening behavior would be accompanied by a decrease in the loss modulus. Clearly, an important focus moving forward should be the inclusion of more sophisticated mechanical models, such as nonlinear viscoelastic models [41, 42], as well as the development of methods for further restricting the acceptable parameter space or for navigating the error surface of the fitting procedures with greater precision, in order to reduce the width of the confidence intervals. Ideally, the methodology should eventually capture all the small features present in the force and indentation data (within noise-related limitations) to maximize the possibility of discovering clear mechanical markers for specific biological conditions.

Despite many mechanical commonalities in cancer cells and tissues, subtle differences may exist in viscoelastic properties that may be critical to understanding tumor progression and metastasis in different situations. It is known that cancerous cells adapt to survive the harsh temporally evolving tumor microenvironment [43], suggesting that many cellular properties, including mechanical properties, must also change accordingly. Several studies have shown that the mechanical properties of the tumor microenvironment play a critical role in cancer progression and metastasis, and have been correlated with the rate of cancer cell migration, proliferation, and resistance to chemotherapeutics [44]. In a tumor, the extracellular matrix is dynamically remodeled, and these modifications create a tumor microenvironment that is stiffer compared with the environment found in normal tissues. This promotes tumorigenesis through downstream signaling, thus forcing the cancer cells to modify their mechanical properties [45]. Previous studies have shown that most cancerous cells are softer than their non-malignant counterparts, and this correlates with their migration properties, whereby softer malignant cells are more migratory [18]. Interestingly, in melanoma cell lines it has been observed that early tumorigenic cells are softer than melanocytes, while highly metastatic melanoma cells were much stiffer than healthy melanocytes [46]. In a developing tumor it is potentially more beneficial to have softer cells initially to allow more deformability and survival in the high pressure intratumoral microenvironment, whereas during metastasis a stiffer cell is more beneficial to enable efficient migration throughout the tight confinements present in tissues. With this complexity in mind, it is reasonable to think that access to more physical parameters such as elasticity and viscosity at multiple timescales should provide much more information to aid the classification of cancer cells during disease progression rather than limiting the analysis to the use of only a deformation-rate-independent stiffness as a disease biomarker.

The potential applications of the method described here are not limited to cancer. Other applications can be conceived, such as, for example, measurement of the viscoelastic properties at multiple timescales in the inner ear sensorial and nonsensorial tissues, in order to elucidate the role of tissue mechanical evolution in hearing loss. Recently, it has been shown that tensional homeostasis provided by non-muscle myosin II contractile activity in the mammalian cochlea is critical for hearing integrity [47]. Inhibition of myosin II contractile activity by Blebbistatin relaxes the organ of Corti and presumably softens it [47]. It has also been shown that TRIOBP activity reorganizes the filamentous cytoskeleton in sensory hair cells and supporting cells, and is implicated in maintaining the mechanical properties of the organ of Corti, while loss of TRIOBP-5 isoform was shown to significantly decrease its apical surface stiffness [48]. Another potential application of our method is the characterization of artery walls with plaque buildups to more deeply understand the role of mechanics in atherosclerosis or other cardiovascular-based diseases. In atherosclerosis, arterial wall increased stiffness is associated with disease pathophysiology and cardiovascular risk events [49]. Notably, these studies and many others still characterize the mechanics of diseased systems by only measuring the elastic properties (stiffness), thus neglecting the rich viscoelastic behavior at different timescales, which as shown here, offers a much more unequivocal path for differentiating cells and tissues.

In summary, we have introduced a novel iterative-fitting method to characterize the viscoelasticity of living cells at multiple timescales based on a simple-to-acquire input consisting of atomic force microscopy quasi-static force curves. The study revealed that unique viscoelastic features emerging at different timescales can be used to precisely differentiate normal from cancerous cells. This approach could thus be exploited to categorize cells for disease diagnosis, monitoring, and treatment. Elucidating the complex viscoelastic behavior of living metastatic cells could also enable engineering of novel therapeutic approaches designed to achieve improved anti-tumor response.

## Materials and Methods

### Cell culture and preparation

Human Foreskin Fibroblast cells were obtained from the American Type Culture Collection (ATCC, Cat #: SCRC-1041, Manassas, VA) and cultured in Dulbecco’s Modified Eagle’s Medium (DMEM, Life Technologies, Carlsbad, CA) supplemented with 10 % fetal bovine serum (FBS, Life Technologies), 1mM sodium pyruvate (Life Technologies), 1x GlutaMAX (Life Technologies), and 1% Penicillin-Streptomycin (Life Technologies). Human Primary Epidermal Melanocyte cells were obtained from ATCC (Cat #: PCS-200-013) and cultured in Dermal Cell Basal Medium (ATCC) supplemented with Phenol Red (ATCC) and Adult Melanocyte Growth Kit (ATCC). Human Melanoma A-375 cells (Cat #: CRL-1619) were obtained from ATCC and cultured in DMEM supplemented with 10% FBS (Life Technologies), 1x GlutaMAX (Life Technologies), 1X Antibiotic-Antimycotic (Life Technologies), and 20mM HEPES pH 7.4.

Cells were plated on glass bottom dishes (Willco Wells, Amsterdam, The Netherlands) to < 70% confluence. Cells were left to adhere on the glass bottom dishes overnight and maintained at 37 °C and 5 % CO_2_. On the following day cells were transported to the AFM system and placed on the AFM X-Y stage to perform force spectroscopy AFM measurements.

### Atomic force microscopy

Live cell measurements were performed using a Bruker BioScope Catalyst AFM system (Bruker, Santa Barbara, CA) mounted on an inverted Axiovert 200M microscope system (Carl Zeiss, Göttingen, Germany) equipped with a Confocal Laser Scanning Microscope 510 META (LSM 510 Meta, Carl Zeiss) and a 40x (0.95 NA, Plan-Apochromat) objective lens (Carl Zeiss). A Petri dish heating stage (Bruker) was used to maintain physiological temperature (37 °C) of cells during measurements. Modified AFM microcantilevers with an attached 25 µm-diameter polystyrene microsphere were obtained from Novascan (Novascan, Ames, IA). The AFM probe spring constant was obtained using the thermal tune method built into the AFM system. Calibrated spring constants for the cantilevers ranged from 0.5 N/m to 1 N/m. After cantilever calibration, the AFM probe was placed on top of the nuclear region of an adherent cell. The deflection setpoint was set between 20 nm and 25 nm, yielding applied forces between 5 nN and 18 nN. The force curve ramp rate was set to 0.5 Hz and the probe speed ranged between 1.9 µm/s and 2.4 µm/s. Multiple consecutive quasi-static force curves were collected on each individual cell with a deflection trigger of 25 nm.

### Data analysis

To evaluate the performance of the new iterative and previous open-search methodologies, a Windows 10 64-bit system with a 9^th^ Generation Intel Core i9 processor and 32 Gb of RAM was used. The Matlab parallel processing toolbox function “parfor” was used to communicate the for-loop orders to each of eight total parallel workers. For every viscoelastic model configuration (i.e., 1-term, 2-term, etc.) 500 fitting attempts were made using the Generalized Maxwell and Generalized Kelvin-Voigt viscoelastic models (Fig. S3) in the Lee and Radok indentation framework (Eq. S19).

For the data analysis of all cell types the Matlab script was modified to function on a High Performance Computing (HPC) cluster at the George Washington University (GWU). For the melanocytes and melanoma, standard computing nodes on the Pegasus HPC (Dell PowerEdge R740 server with Dual 20-core 3.70 GHz Intel Xeon Gold 6148 processors, 192 GB of 2666 MHz DDR4 ECC Register DRAM, 800 GB of onboard SSD storage, and Mellanox EDR Infiniband controllers to 100 GB fabric) were utilized to perform the same parallelized iterative fitting methodology and the augmented hardware specifications allowed 20 parallel workers to be used for each dataset. The increased processing power enabled a larger number of fitting attempts for each dataset. Specifically, 1000 fitting attempts based on random starting points were used for every model configuration. The processing time was dependent upon the length of the dataset under study but varied between 4 and 17 hours. For the fibroblasts, due to time and computing cluster availability constraints at GWU, the National Institutes of Health (NIH) Biowulf cluster was used with a similar code repository (modified to meet specific HPC requirements for job scheduling). The 28-core “norm” queue for Biowulf (Dual 28-core x 2.4 GHz Intel E5-2680v4 processor, 256 GB of RAM, and a 56 Gb/s FDR Infiniband controller) was used for fitting, which allowed a maximum of 28 parallel workers. Due to the large number of cells included (33), the fibroblasts required approximately 147 hours of total runtime on Biowulf.

The viscoelastic results presented in this manuscript were calculated using the optimal parameter set obtained from analyzing the average of 70, 71, and 193 force curves from the melanoma, melanocyte, and fibroblast cell conditions, respectively. The averaging was performed in the frequency domain, after each optimal parameter set was used to calculate a predicted frequency-dependent curve for the storage modulus, loss modulus, and loss angle from every force curve. The solid lines in Fig. 4 and Fig. 6 are the simple average of these individual viscoelastic harmonic quantity predictions, and the confidence bands (shaded regions) are the range of observations for each discrete frequency using a 95% confidence level and the Student’s t distribution. In total, the averaged datasets contain curves collected from 13 unique melanoma cells, 12 unique melanocytes, and 33 unique fibroblasts.

The Matlab script used the “lsqcurvefit” (least squares) function with the “trust-region-reflective” internal algorithm selected. The cost function used was a simple Sum of the Squared Error (SSE) between the integral term in Supplemental Information Eq. S4 and Eq. S5, and the normalized y-data (left-hand side) for both equations, respectively. The lsqcurvefit() function was chosen over other alternate nonlinear least-squares fitting functions because it showcased an increased speed of convergence compared to other options (specifically fmincon()), and the trust-region-reflective algorithm was used because it is the default for that function.

The code used to acquire the optimal parameter sets obtained in this manuscript is available in a github repository [50]. The datasets for each cell type (melanocytes, melanoma, and fibroblasts) have also been included in that repository, in the “data” directory.

## Acknowledgments

S.D.S. gratefully acknowledges support from the U.S. National Science Foundation, through award No. CMMI-2019507. A.X.C.R. acknowledges support from the National Institutes of Health (NIH) Intramural Research Program fund of the National Institute of Biomedical Imaging and Bioengineering (NIBIB; grant # ZIA EB000094) and the NIH Distinguished Scholars Program.

## Supporting Information

### 1. Phase Contrast Light Microscopy Images for 2D Adherent Human Metastatic Melanoma Cells and their Normal Counterparts

**Fig. S1.**
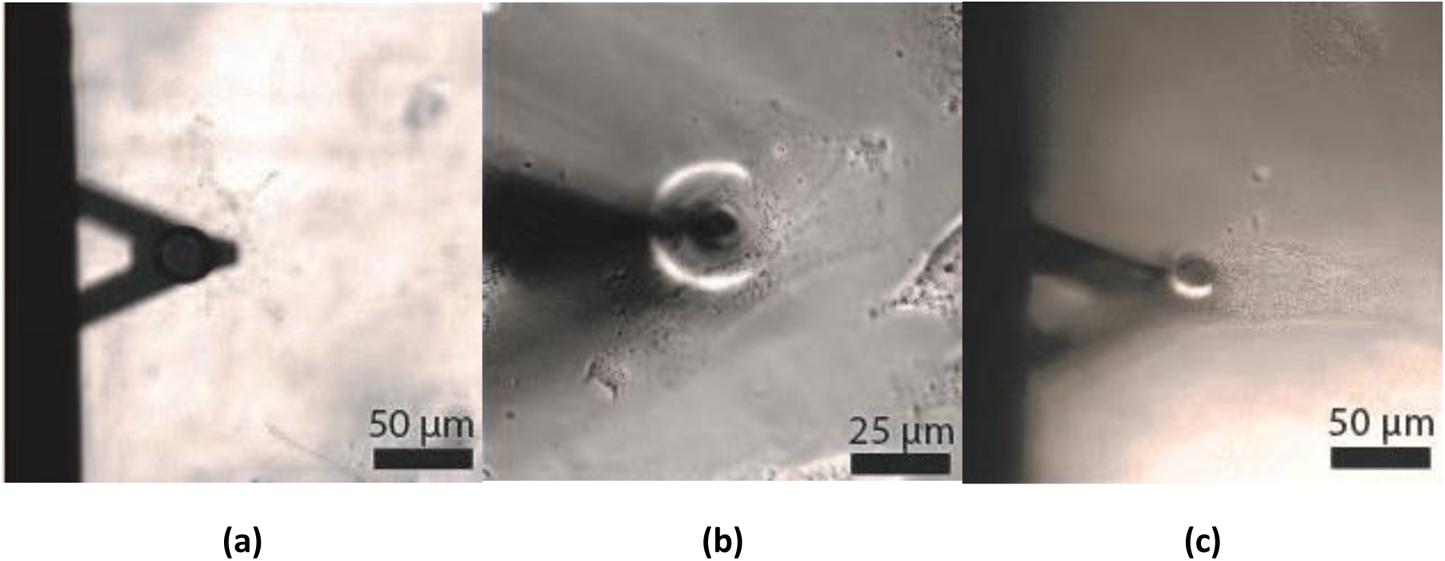
Phase Contrast Light Microscopy Images for Each Adherent Skin Cell Type under Study. This figure shows phase contrast light microscopy images for a 2D adherent human primary epidermal melanocyte cell (a), human metastatic melanoma A375 cell (b), and human foreskin fibroblast (c).

### 2. The Lee and Radok Framework for Spherical Indentation of a Viscoelastic Half-Space

To cast the problem of a spherical probe indenting a viscoelastic surface, it is necessary to make a series of simplifying assumptions about the tip-sample interaction. The longstanding geometry which Lee and Radok [1] proposed in 1960 applies for spherical, rigid indenters forced into a viscoelastic half-space, and is visualized in Fig. S2.

**Fig. S2.**
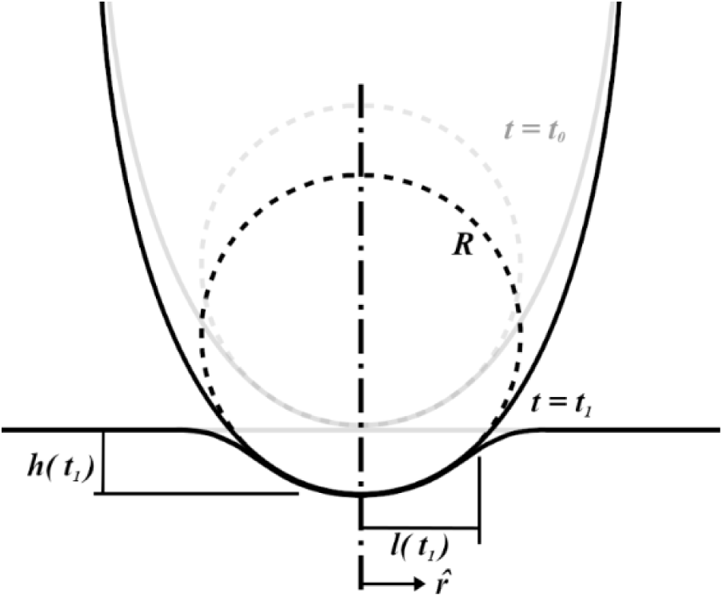
Spherical Indentation Geometry Proposed by Lee and Radok [1]. The deepest indentation occurs at the sphere’s central axis (r = 0) and is labeled as h(t); the indenter has a radius of curvature R; the distance from the central axis to the edge of contact is labeled as l(t) and is also commonly referred to as a(t).

A detailed breakdown of the derivation and associated assumptions for this framework has been provided previously [2], but it remains critical for users to confirm the validity of this approach for their specific conditions before implementing it. To that end, at least the following conditions must hold for the Lee and Radok derivation to be applicable:

1. The tip shape can be regarded as approximately spherical;
2. The indentation depth is sufficiently small, such that the strain behaves approximately linear;
3. The contact area is monotonically increasing during the experiment;
4. The problem can be cast as quasi-static (time is only a reference variable).

If the above can be assumed, the following relationship between applied load (*F*) and indentation depth (ℎ) applies for a viscoelastic material having Poisson’s ratio ν (commonly assumed to be incompressible, such that ν = 0.5), and characteristic operators *u* and *q*:

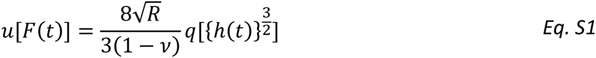

These characteristic operators are described as a polynomial series in the complex Laplace domain variable “s”, and are related to the material Retardance (*U*(*s*)) and Relaxance (*Q*(*s*)) by:

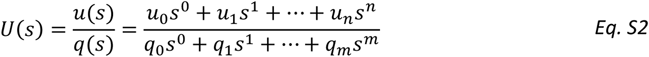

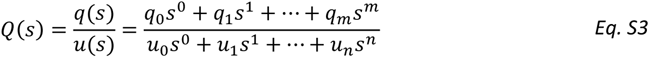

The Lee and Radok technique is particularly powerful for simulating tip-sample interactions in atomic force microscopy (AFM) because differential equations in the time domain can be cast as simple polynomials in the Laplace domain, provided that the viscoelastic model can be successfully described and transformed into the Laplace domain. Relatively straightforward rearrangement of *Eq. S2* and *Eq. S3* allows each order of the variable *s* to be grouped, and the coefficients for each series can then be calculated using known material parameters. If either the force or indentation is provided, and the Lee and Radok assumptions still hold, calculating the material response simply requires taking discrete derivatives of the AFM observables.

Since the material models have been described using *U* and *Q*, *Eq. S1* is transformed into the Laplace domain and rearranged such that the indentation is the output variable. Thus:

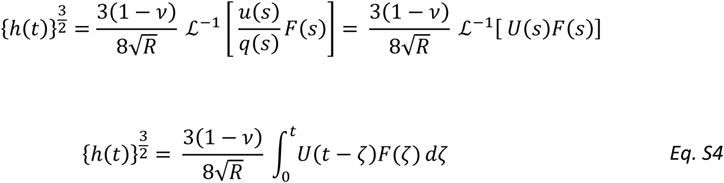

Here, the retardance of the Generalized Kelvin-Voigt model can replace the Retardance (*U*) in *Eq. S4*. Since the force and indentation are known throughout an AFM tip-sample interaction, the only unknown parameters are the compliances and characteristic times contained within *U*. This equation can also be easily re-written to utilize the Relaxance (*Q*) of the Generalized Maxwell Model:

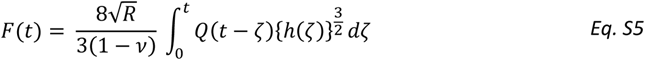

### 3. Guidelines for the Choice of a Viscoelastic Material Model

For many problems in viscoelasticity, it is standard to assume that various combinations of springs and dashpots in series and/or parallel can reproduce the viscous and elastic material behavior observed during mechanical interactions. These descriptions are commonly known as “rheological” or “spring-dashpot” models, and usually fall into the broad category of “linear viscoelasticity”—a term used for models that assume the elastic and viscous action of a material can be analytically separated. These models are inherently limited in scope to small stresses or indentations and must be assembled into more complex configurations to successfully reproduce commonly observed material behaviors such as strain creep (increasing strain under applied constant stress) and stress relaxation (reduction in stress under applied constant strain). Intermediate combinations, such as the Standard Linear Solid (SLS) or Burgers models, still contain relatively few parameters for describing samples but can be useful for initial approximations or temporally limited datasets. For further discussion of these models, including the strengths and weaknesses of each, the reader is directed to the available literature [3, 4].

Recently, Bonfanti et al. published an introduction on soft matter viscoelasticity, with emphasis on fractional power law models [5]. There also exist manuscripts by Efremov et al. which treat viscoelasticity in soft biological samples examined using AFM techniques, with an emphasis on parameter extraction [6] and on reviewing models applied in the literature [7]. However, all these publications focus little on utilizing models with a *discrete* number of retardation times to reproduce material behavior, in favor of power law models or the simpler intermediate combinations mentioned above. The benefit of fractional models, and other forms of continuous relaxation spectra approaches, is the lack of transition zones in the material response and the relatively small number of model elements required to describe material action across many timescales. While this is convenient for obtaining material property approximations with fewer parameters (thus reducing computational overhead and leading to simpler representations), the shape of the relaxation spectra is dependent upon the number of terms. This means that if there is action on a timescale deemed interesting, it is difficult to separate and monitor that term because the power law model parameters control the shape of the response for the entire timescale spectrum. The issue is further exacerbated when many distinct features are present in a mechanical experiment dataset. In such a case, having discrete retardation times provides a distinct advantage, as one can observe the relative shift in characteristic time and stiffness of an element, which may correspond to specific markers in the harmonic response of the material, and differentiate between specific materials or, for example, biological material conditions, by monitoring changes in well-understood harmonic quantities such as the storage and loss moduli.

When dealing with cells, specifically, it is necessary to forego convenient, simplistic material models and provide a sufficiently large parameter set, such that complex material action linked to important timescales can be identified. The most straightforward implementation is a generalized viscoelastic model, with a dynamic (i.e., user-defined) number of discrete elements. As such, the critical contribution from this manuscript is a detailed outline of an iterative approach, which can provide important feedback for the user, and help to avoid both “overfitting” and improper extrapolation of the material response beyond reasonable limits. This methodology represents a significant step toward accurately characterizing materials where action can be linked to specific timescales and is necessary to support biological research efforts seeking new mechanical markers in cells.

**Fig. S3.**
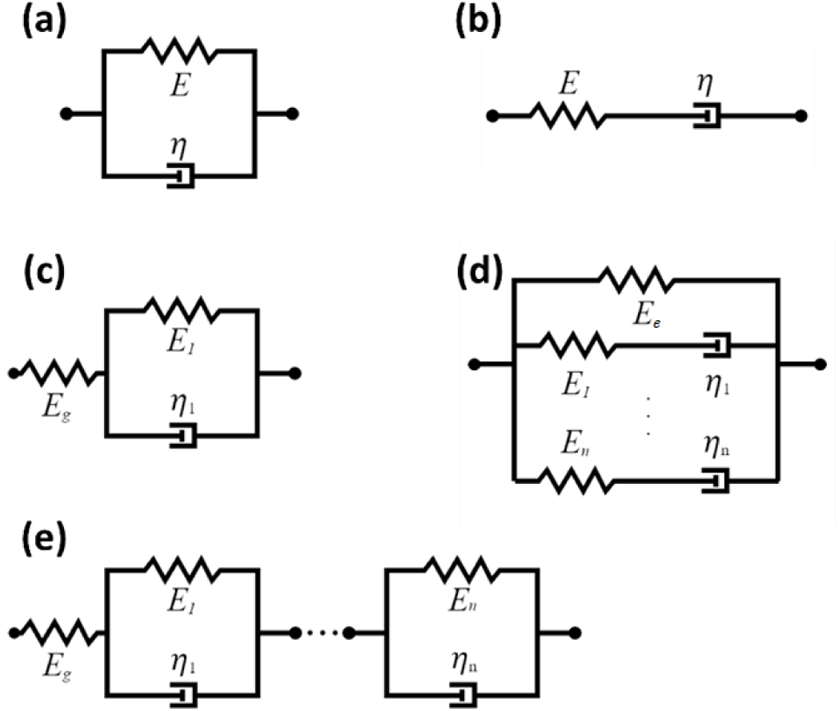
Common Viscoelastic Models. Viscoelastic descriptions are fundamentally comprised of spring (elastic) and damper (viscous dissipative) elements; (a) and (b) are the Kelvin-Voigt and Maxwell elements respectively; (c) is the Standard Linear Solid (SLS) in the Kelvin-Voigt configuration; lastly, (d) and (e) are the generalized models discussed in this manuscript, the Generalized Maxwell and Generalized Kelvin-Voigt, respectively. Note that the SLS model can also be configured to use a Maxwell element, wherein (b) is placed in parallel with a spring. As is discussed in this appendix, the Maxwell and Kelvin-Voigt models are mechanically equivalent and can be interconverted, although the conversion is not always simple when handling the generalized forms (c & e).

The Generalized Kelvin-Voigt and Generalized Maxwell models are illustrated in Fig. S3, along with their sub-components. Since they are conjugate models, they can both represent the same range of material action provided their Prony series are correctly parameterized [8]. However, it is commonly understood that the suitability of each model is dependent upon the problem boundary conditions. The Generalized Maxwell model is most easily described in terms of Relaxance (the material stress response to applied strain), whereas the Generalized Kelvin-Voigt model is conveniently formulated in terms of Retardance (strain response to applied stress) [3]. Approximate analytical relationships and numerical interconversion methods have been introduced to translate between the Prony series parameters of the two models [9].

These formulations can be considered the viscoelastic-counterparts to the Young’s Modulus and Glassy Compliance of linear-elastic models. Whether elastic or viscoelastic, these quantities act as conversion factors between stress and strain. Unlike for the linear-elastic case where these operators are constant values, the viscoelastic Relaxance and Retardance are time-dependent.

Full derivations of the constitutive equations for each model pictured in Fig. S3 are available in the literature, which are based on the Laplace Transform [3, 8] and fundamentals of continuum mechanics [10], but are beyond the scope of this appendix. As such, we begin by describing the viscoelastic Relaxance (*Q*) and Retardance (*U*) in terms of stress (σ) and strain (ε):

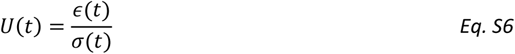

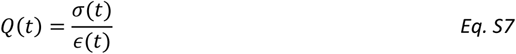

These relationships are general and will need to be specified to the AFM static force spectroscopy (SFS) problem geometry later using one of the available indentation frameworks. For now, it will suffice to provide descriptions of each model using their appropriate boundary conditions. The Laplace Domain treatment of the mechanical descriptions in Fig. S3 is the simplest method for deriving relationships for *U* and *Q*, using a technique analogous to the Electrical Impedance Method in circuit theory. Springs and dashpots are combined using common series and parallel rules to provide a single-element description—similar to a transfer function. As shown by Tschoegl [3], the Generalized Kelvin-Voigt model features the following Retardance

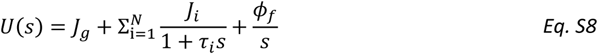

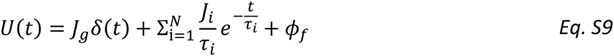

This description assumes that the model contains both a Glassy Compliance (*J_g_*) and a single steady-state fluidity (ϕ_f_) in addition to N Voigt elements in series, each with their own compliance (*J_i_*) and characteristic time (τ_i_). The Relaxance of a Generalized Maxwell solid can be similarly described as:

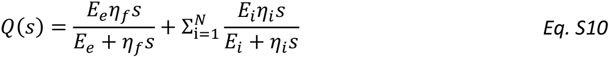

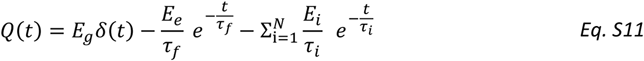

Here, the Elastic Modulus ( *E_e_*), Glassy Modulus (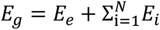), and steady state viscosity (η_f_) have been included in addition to the moduli (*E*_i_) and characteristic times (τ_i_) of each Maxwell element up to N branches. Note that the damper viscosities have been replaced by the element characteristic times according to 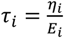 in *Eq. S11*. Additionally, in the case that the_*E*i_ material is arrheodictic (i.e., it is unable to sustain steady-state flow), the second term in *Q*(*t*) is excluded. This eliminates any relaxation of the model arm containing *E_e_*, meaning that as time trends toward infinity the value of *Q*(*t*) will approach *E_e_*; conversely, if the material is known to be rheodictic, then the term is included and the Relaxance will trend toward zero at long timescales. A lack of steady-state fluidity in the Generalized Kelvin-Voigt model indicates that the final term in the series *U*(*s*) is removed. The time-dependent nature of these relationships means that for each calculation the entire stress or strain history must be known, then convolved with the appropriate spectrum to recover its pair.

It is worthwhile to note that the existence of an elastic term and steady-state fluidity in the material is not always reasonable. Lopez et al. suggested that the fluidity could be neglected for shorter indentation experiments (on polymers), as the overall interaction time would be well under the threshold for shear flow [8]. While true for many polymers, biological samples are often much softer and more fluidic at standard operating temperatures and pressures. As such it is recommended to include at least a single steady-state fluidity term when considering biological samples, with the understanding that more complex treatments may be necessary, such as the inclusion of fluidities at multiple timescales. Similarly, the elastic term may be negligible for highly fluidic samples. For the Generalized Maxwell case, the idea that a static stiffness acts in parallel with the other elements may be violated when the samples are sufficiently soft, and a clear majority of the energy is dissipated by viscous action. If an elastic term is included under such conditions, one could erroneously attribute the stiffness of short timescales to the elastic term and the total dissipated energy would be underestimated, because the elastic energy is returned after unloading the sample. It should be noted that, for the adherent skin cell lines studied in the accompanying main manuscript, the steady-state fluidity was excluded for simplicity. The performance of the methodology on cells, while also utilizing new configurations of the generalized viscoelastic models in rheodictic form, remains to be investigated.

There are several critical points to keep in mind when applying the parameterization methods presented. First and foremost, the methodology does not naturally prevent “overfitting” a dataset; in this context, overfitting occurs when a user inserts a larger number of model terms than is necessary to acquire a closer approximation to the data. Iteratively introducing terms of increasing timescale gives a clear indication of when the minimum number of terms necessary for a close estimate occurs, but users must be careful to include only orders of magnitude that can reasonably be justified as being truly present in the dataset. For example, the likelihood that an indentation occurring for 1 millisecond contains mechanical artifacts from timescales on the order of 10^0^ is low and including a characteristic time on that order could obscure the effects of lower order terms. To avoid this issue, the smallest number of terms possible that provide an adequate fit (as defined by the user) should be chosen, and iteratively introducing terms clearly indicates when that occurs.

In addition to controlling the overall number of terms, the upper and lower bounds for all parameters must be appropriately chosen to avoid poor fitting performance and violating the physical principles of the material. For example, fitting a Generalized Kelvin-Voigt model without restricting the individual compliances of each term to values above zero would allow terms to apply negative (downward) forces from within the material, which would be unreasonable for a repulsive indentation experiment (where the material resists the tip motion). While this case is clear for the stiffness terms, the characteristic times are slightly more complex to consider. It is not unreasonable to specify specific values for each characteristic time and allow the stiffnesses to vary within the bounds discussed above, although this approach makes an implicit assumption that those characteristic times exist within the material. In this case, the user has limited the parameter space and may be able to acquire results quickly and with good accuracy. However, by removing the characteristic times as a degree of freedom in the fitting, one may obscure valuable insight into the sample’s mechanical nature. On the contrary, by allowing the characteristic times for each term to vary within some orders of magnitude, users leave open the possibility that a more natural combination of parameters can represent the material to a higher degree of accuracy at the cost of computational overhead. When action, whether it be hardening, softening, or otherwise, is known a-priori to occur within a sample on a specific timescale, users should consider including specific terms in the model description. Inevitably, the purpose of these parameterization efforts is to acquire a deeper understanding of the mechanical response of the sample and failing to leverage previous knowledge can lead to unnecessary additional time and effort.

In extrapolating material behavior using parameterized models, it is critical to acknowledge the limitations of the parameters obtained by fitting a discrete dataset. The sampling frequency determines the lower (shorter) bounds of the window within which the model parameters are useful—action on timescales smaller than the sampling frequency cannot be accurately resolved within the dataset, and as such, it is unknown whether the material action will be properly predicted for a different set of experimental conditions where tip velocities are significantly faster. For instance, an SFS-based parameter set would be difficult to use for estimating tapping-mode AFM indentations because the interaction times in the latter technique are orders of magnitude shorter. Furthermore, at smaller timescales the sensor noise floor could play a dominating role in the “perceived” forces, leading to large errors. Similarly, the total repulsive indentation time determines the upper bounds of applicability for the model parameters. It would be inappropriate to assume that mechanical action applied over timescales that are several orders of magnitude larger than the duration of force application can be adequately discerned from the smaller-timescale response measured. For capturing data during long experiments, tightly controlling external sources of noise and error (room temperature, humidity, vibration, instrument drift, etc.) would also become increasingly important, as these changes could erroneously materialize in the dataset as stress-induced material action.

It is worthwhile to discuss the quantities of merit that can be equated between unique experiments and samples. Specifically, leveraging individual terms within a parameterized series is an invalid form of comparison. This is because there exist a vast number of theoretically appropriate parameter sets that can recreate the mechanical action contained within a single indentation dataset. To illustrate this point, best-fit parameter sets can contain individual stiffnesses that appear to vary widely, disproportionate with the overall rigidity of the sample. Until a method for differentiating between valid and invalid optimal parameter sets can be developed, it remains inadvisable to utilize specific model terms independently. Using the predicted stiffness of all terms within the model together is more appropriate, as the fitting algorithm optimizes the entire series simultaneously to reproduce the apparent force-indentation relationship. The storage modulus, loss modulus, loss angle, and retardance (or relaxance) provide the current best measures for comparison between individual experiments. Calculation of the viscoelastic harmonic quantities from a parameterized model has been covered in several previous publications [8, 2].

Lastly, and most fundamentally, the assumptions presented during the framework derivations must remain valid throughout the duration of the experiments. In particular, the linearity assumptions dictate that the indentation depth must be “sufficiently small” such that the force curves appear approximately linear. While acceptable for hard materials and fast indentations, this condition could be easily violated for soft or biological samples. There is no specific numerical threshold for an indentation being too deep, although it is common to observe the indentation strain measure 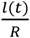 and treat it similarly to the engineering strain quantity 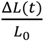. The commonly referenced (though flexibly defined) transition from “linear strains” to “finite strains” then becomes applicable for the indentation frameworks presented, and strain values under 1-2% would normally ensure the linearity requirement for many common materials, depending on the material.

### 4. Interconversion between the Generalized Kelvin-Voigt and Generalized Maxwell Models

It is known that the Generalized Maxwell and Generalized Kelvin-Voigt models are “mechanically equivalent” to one another. Analytically this is enforced by the definitions given above (*Eq. S2* and *Eq. S3*), which reveal that the Relaxance (*Q*(*s*)) and Retardance (*U*(*s*)) are inverses of one another in Laplace space. Since the Generalized Maxwell and Generalized Kelvin-Voigt models can be written in terms of Relaxance and Retardance, given that the observed material action is the same in both cases, there must exist a set of parameters that satisfy the simple relation *Q*(*s*)*U*(*s*) = 1. This understanding is critical when experimental convenience demands implementation of one model over the other—the need for well-behaved system excitations (e.g., step-inputs) in stress or strain can require users to utilize a stiffness- or compliance-based model. In such situations, the user must understand that an appropriate parameterization of one model, given that it reproduces the input datasets to a high degree, will supply the parameters for its pair through one of several methods:

1. Using the extracted parameters to “simulate” the response to a step function in stress or strain using *Eq. S4* or *Eq. S5* and fitting the desired model to this new dataset;
2. If the models are simple enough (e.g., if they contain only a single element, such as for the SLS), analytical relationships exist for converting between parameters [3];
3. The Collocation Method, which involves calculating the complex s-domain distribution of one model and fitting the other model’s parameters to the result.

If expediency is important, option 1 is convenient because it does not require conversion to the Laplace domain for either model (although this step is analytically straightforward). In many cases, option 2 is not feasible—beyond a single element, the relationship between models involves fourth order polynomials in the parameters and is ill-posed. Even for two elements, the mathematical treatment required would easily exceed the time or patience of most analysts.

The remaining option is to use the s-domain Collocation Method outlined by Tschoegl [3], which involves using the Laplace Domain forms of the Relaxance and Retardance (*Eq. S8* and *Eq. S10*) with the understanding that *Q*(*s*)*U*(*s*) = 1. As such, evaluating *Q* or *U* for a wide range of s-values will show a shape similar to an inverted-logistic function with several features in the transition region. The user can then perform a fit using the inverse of the complementary spectra to obtain the parameters that create mechanical equivalence between the two models.

Finally, if the representation of a model in the harmonic space is desired, one need not perform a fit to obtain equivalent parameters before calculating the relevant harmonic quantities (loss modulus, loss compliance, storage modulus, storage compliance, loss angle). Instead, relationships exist that allow for conversion between these quantities, based on the use of the absolute modulus (*Ẽ*) and absolute compliance (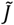):

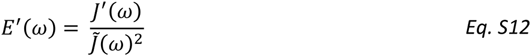

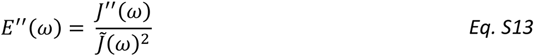

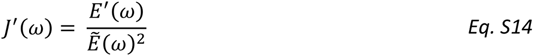

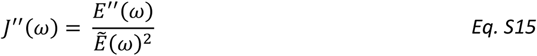

Where the absolute modulus and compliance are calculated as:

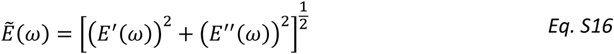

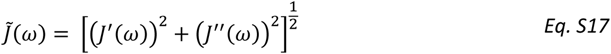

Thus, if the desired output quantities are the loss modulus (*E*′′) and storage modulus (*E*′) when the user has a parameterized Generalized Kelvin-Voigt model (i.e., with native loss compliance *J*′′ and storage compliance *J*′), these relationships can be used to convert between the quantities successfully. This is possible because these distributions exist in the complex domain—conversion between time-domain quantities still requires fitting (e.g., *Q*(*t*) to *U*(*t*)). For a full suite of the equations necessary to calculate the harmonic moduli, the reader is directed to the literature [3, 8, 2].

### 5. Summary of Equations Required for Implementation of the Methodology

With this understanding of the generalized models and indentation framework one can begin extracting parameters from properly conditioned AFM SFS datasets. The steps required for conditioning raw AFM observables for fitting have been previously enumerated [2] and are thus excluded here. The following equations are the only ones necessary to extract material information using the generalized viscoelastic models and the parameter extraction methodology:

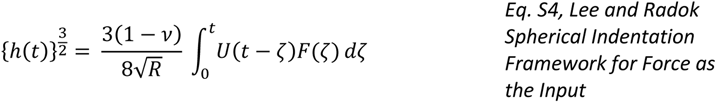

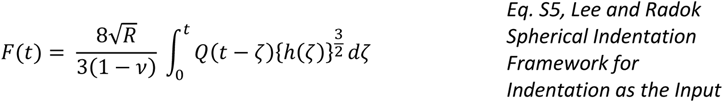

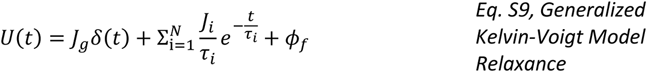

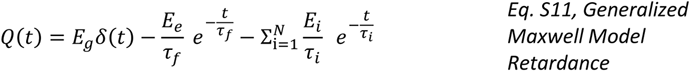

### 6. Data Simulation Methods

To adequately characterize the performance of both iterative- and open-search methods, AFM SFS data was simulated using known viscoelastic parameters and a dynamically-generated differential equation series. Because the Lee and Radok method only applies rigorously for monotonically increasing contact area, only the approach portion of the SFS curve was simulated. As previously stated, the Laplace transform and viscoelastic operators *U* and *Q* can be used in conjunction with a viscoelastic model to create a polynomial representation of the material. By using an approach similar to the calculation of electrical impedances in a circuit, the material descriptions from Fig. S3 were reduced to Eq. S8 and Eq. S10. At this point, the series can be easily rearranged to have the same form as Eq. S2 and Eq. S3, using standard algebraic operations. However, as the number of terms included in the viscoelastic description increases, these polynomials become increasingly complex. Beyond two or three additional elements, the fourth order polynomials are onerous to derive manually. In this case, utilizing a symbolic computational toolbox (such as the Matlab Symbolic Toolbox, Wolfram Mathematica, or Python’s SymPy) can more readily handle the necessary simplification, factorization, and re-organization. Providing Eq. S8 and Eq. S10 to a symbolic programming tool can thus dynamically generate the coefficients required to simulate the following well-known differential representation of viscoelastic materials:

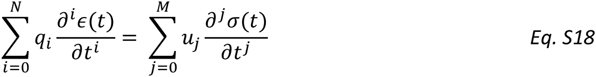

Here the variables M and N represent the highest order of s associated with the desired polynomial series, and are denoted as separate exclusively for the case where u(s) and q(s) do not contain the same order of s. Importantly, when the Generalized Kelvin-Voigt representation utilizes a steady-state fluidity term, the polynomial will also feature an order of s^−1^, which in the time domain represents integration from the onset of indentation to the current time. The stress and strain in *Eq. S18* are then replaced by functions of force and indentation. The following relationship can be used to generate AFM force curve data for the Lee and Radok (*Eq. S19*) indentation framework:

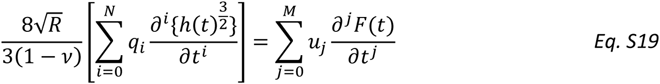

Once the coefficients *u*_i_ and *q*_i_ are available, one can simulate the tip motion in time by programmatically beginning with the AFM probe located at an initial height, and the enforcing an approach velocity towards the surface for the cantilever base. Each timestep requires solving one or more standard kinematic equations of motion for the AFM cantilever, depending on the number of normal modes included in the cantilever model. In addition, discrete derivatives to the N^th^ order are necessary at each timestep, such that *Eq. S18* can be rewritten to solve for the highest derivative of the force. Using the strain and strain rates for the current timestep, in addition to the force and force rates (below the highest order) from the previous iteration, one can calculate the highest derivative of the force for the current time. This value can then be discretely integrated to provide all force terms at the current time, including the lowest-order term corresponding to the tip-sample force. The resulting force can be added to any long-range tip-sample force model (e.g., to account for nonbonded forces such as van der Waals attraction, etc.) and fed forward to the next timestep as input into the tip acceleration. The selected timestep must be small enough to ensure stability of the simulation. Using a timestep on the order of 10^−10^ seconds for the tip-sample contact portion of the force curve is small enough to prevent divergence in most cases. If the user wishes to adjust the timestep for their own simulation, taking the period of their cantilever’s primary mode (the inverse of the cantilever natural frequency) and selecting a timestep several orders of magnitude lower (at least 4) is an adequate starting point. For these simulations, the intended outcome was an evaluation of the computation time, memory requirements, and accuracy of the methodology.

The cantilever properties used for the simulation were chosen to match a compliant silicon-cantilever colloidal probe. The tip radius was chosen to be 12.5 µm, with the tip attached to a cantilever of stiffness 0.6 N/m, with a fundamental frequency of 75 kHz and quality factor of 2. The sample was approached at 2 µm/s, starting from 1 µm away. The long-range tip-sample forces were not included, and indentation continued until a trigger force of 5 nN was achieved. The surface was given an elastic modulus of 9220 Pa, a Poisson’s Ratio of 0.5, and steady-state fluidity was excluded. The Prony series parameters used are provided in Table S1.

**Table S1.** Viscoelastic Prony Series Parameters Used for Data Simulation

| Model Element Number | Stiffness (Pa) | Characteristic Time (s) |
| --- | --- | --- |
| 1 | 6691 | 6.84E-4 |
| 2 | 1877 | 4.40E-3 |
| 3 | 9684 | 2.97E-4 |

### 7. Comments on Convergence and Benchmarking

For a critical portion of the process, the outlined viscoelastic parameter extraction methodology relies heavily upon accurate numerical approximations of the data. As such, the issues encountered cease to be analytically centered, and instead involve the programming implementation of those relations. Determining “convergence” and establishing well-behaved benchmarks for performance are important steps to validating that the implementation does, indeed, provide the insight that is claimed by the user.

On the subject of convergence, there are several quantities that merit discussion. First and foremost is the number if fitting attempts made. This is defined here as the number of unique iterations where the algorithm of choice is given the model description, output data, and a starting point for each parameter. To converge to a global minimum error with the current methodology, it is necessary to attempt a sufficiently large number of unique starting parameter combinations, such that the parameter space is widely explored. This clearly incurs computational overhead but is necessary because the final parameters are very sensitive to their initialization values. If the number of iterations is too low, then the likelihood is diminished that the model was provided enough opportunity to achieve good convergence. Instead, the number of iterations must be sufficiently high that parameter sets on their way to high-fidelity approximations can fully converge to a solution. However, allowing too many iterations can also lead to poor performance, especially when the optimization algorithm encounters a plateau in the parameter space. Such plateau regions can be large and may have no discernable negative gradients. In this case the algorithm will take small steps and simply run out of iterations without making any significant progress. If the allowed number of iterations is too large the algorithm can waste considerable amounts of time attempting to converge a poorly conditioned parameter set.

Benchmarking the performance of the programming implementation usually requires processing known (or well characterized) datasets and evaluating the fitting quality and resources used against an established approach. In the case of viscoelastic parameter extraction, this would usually involve using a simulated dataset with known parameters to which the methodology is applied. The user can then assess how well the results match the known parameters of the model and decide whether that level of accuracy is enough for their application. It can also be beneficial to simulate datasets that resemble the expected experimental data in both timescale and order of magnitude. This allows a priori optimization of the number of iterations, fitting attempts and range of randomized input parameters. Clearly, measurement noise and other factors will influence the process when using real data, but selecting a tighter range of randomized starting points and optimizing the parameter choices can speed up convergence.

### 8. Comments on Common Troubleshooting Methods

Most often, performance issues can be identified under a few categories (in order of decreasing frequency, in our experience):

1. Settings for the fitting attempt (not allowing enough iterations, using a poorly chosen range for initial guesses, etc.);
2. Outliers in the dataset (e.g., when averaging curves, for example, keeping one or more datasets which showcase highly erratic material action due to measurement instabilities);
3. Coding implementation (programming bugs, introduction of error due to discrete approximations, poorly chosen algorithms, functions with default settings that cause confusion, etc.);
4. Analytical implementation (issues with the derivation of the equations as implemented in the code).

Of all these categories, the coding implementation errors can be the most time consuming to fix. It usually is not immediately clear when a coding error is causing the problem, which reinforces the need to run common benchmarking operations *before* evaluating real data. Provided the simulated data is well understood, this can showcase programming issues more quickly. Building in simple-to-activate feedback (e.g., plots, status messages, etc.) can also speed up this process, at the expense of initial complexity.

If changing the number of allowed iterations or fit attempts does not rectify the issue under consideration, is necessary to begin by confirming that the viscoelastic models appear to act as expected in the code. Often, small issues like referencing the viscoelastic parameters in an incorrect order can cause confusing intermediate outputs which may not be immediately obvious. Checking that any subfunctions are working properly, are up-to-date, and any old versions are deleted or out of memory usually follows.

Lastly, and anecdotally frequent, are hidden settings in standard functions. For example: the convolution functions in Matlab and Python generally default to a “full” setting. This means that the output vector from the convolution of two vectors with length N and M respectively will be N+M-1 in length. If this is not taken into account, inappropriate clipping of the data may occur. Behaviors of functions that are unexpected, or worse, *change* with new distributions of the coding environment, can be difficult-to-identify sources of error. As before, it remains important to build in safeguards that notify the user when data is ill-conditioned or unexpected data manipulations are being performed.

